# Functional Changes to Achilles Tendon and Enthesis in a Mouse Model of an Adolescent Masculine Gender-Affirming Hormone Treatment

**DOI:** 10.1101/2024.06.10.598308

**Authors:** LeeAnn A. Hold, Tessa Phillips, Paige Cordts, Steph Steltzer, Seung-Ho Bae, Brandon Henry, Nicole Migotsky, Sydney Grossman, Cynthia Dela Cruz, Vasantha Padmanabhan, Molly Moravek, Ariella Shikanov, Adam C. Abraham, Megan L. Killian

**Affiliations:** Department of Molecular and Integrative Physiology, Michigan Medicine, Ann Arbor Michigan, United States; Department of Orthopedic Surgery, Michigan Medicine, Ann Arbor, Michigan, United States; University of Toledo College of Medicine, Toledo, Ohio, United States; Michigan State College of Osteopathic Medicine, East Lansing, MI, United States; Department of Biomedical Engineering, Michigan Medicine, Ann Arbor, Michigan, United States; Department of Obstetrics and Gynecology, Michigan Medicine, Ann Arbor, Michigan, United States

**Keywords:** Biomechanics, Enthesis, Estrogen, Tendon, Testosterone

## Abstract

Many transgender youth seek gender affirming care, such as puberty suppression, to prolong decision-making and to align their physical sex characteristics with their gender identity. During peripubertal growth, connective tissues such as tendon rapidly adapt to applied mechanical loads (e.g., exercise) yet if and how tendon adaptation is influenced by sex and gender affirming hormone therapy during growth remains unknown. The goal of this study was to understand the how pubertal suppression influences the structural and functional properties of the Achilles tendon using an established mouse model of transmasculine gender affirming hormone therapy. C57BL/6N female-born mice were assigned to experimental groups to mimic gender-affirming hormone therapy in human adolescents, and treatment was initiated prior to the onset of puberty (at postnatal day 26, P26). Experimental groups included controls and mice serially treated with gonadotropin release hormone analogue (GnRHa), delayed Testosterone (T), or GnRHa followed by T. We found that puberty suppression using GnRHa, with and without T, improved the overall tendon load capacity in female-born mice. Treatment with T resulted in an increase in the maximum load that tendon can withstand before failure. Additionally, we found that GnRHa, but not T, treatment resulted in a significant increase in cell density at the Achilles enthesis.

**NEW & NOTEWORTHY:** These findings demonstrate that puberty suppression or testosterone does not negatively influence tendon structural or functional properties in a mouse model of transmasculine gender affirming care. In all treatment groups, the ability of the tendon to withstand load was significantly increased. Puberty suppression with GnRHa significantly increased enthesis cell density, suggesting an extended growth phase. These findings elucidate the effects of gender affirming care on the structural and functional properties of the tendon and enthesis.

## INTRODUCTION

Sex differences have been associated with increased risk of musculoskeletal soft tissue injury, especially for the female-born patient (1). The risk of injury is especially elevated in female-born athletes, who have a 4-8-fold increase in risks associated with anterior cruciate ligament tears (ACL) compared to male-born athletes (2, 3). This risk is proposed to be enhanced during the follicular and ovulatory phase of menstruation, when estrogen is peaking, as a prospective study of the rate of ACL tears in female-born skiers showed tears were 2.4 times more frequent during this time (2). Estrogens have been identified to be a regulating factor in many musculoskeletal tissues, including tendon (4–6). Tendons express estrogen receptors, and estrogen plays a critical role in collagen synthesis in tendon (7). For example, estrogen deficiency in rats following ovariectomy can lead to ∼30% reduction in collagen content in the Achilles tendon (8). Testosterone contributes to increased tendon stiffness in male-born individuals by increasing collagen turnover and content (1). Furthermore, testosterone may indirectly reduce tendon and ligament laxity by downregulating the expression of relaxin receptors, a modulator for joint elasticity (9). While these studies show a role of sex steroids in mature tendon composition, adaptation and injury, the impact of sex steroids on tendon function during pubertal growth has not yet been explored.

Transgender and gender diverse youth seek gender affirming care to align their physical sex characteristics with their gender identity, which is important and beneficial for the overall mental health and wellness of the individual (10). One approach to gender affirming hormonal therapy (GAHT) is the use of gonadotropin release hormone analogue (GnRHa) to prevent further development of the endogenous secondary sex characteristics corresponding to the individual’s sex designated at birth (11). Gonadotropin release hormone (GnRH) is released by the hypothalamus, stimulating the release of pituitary gonadotropins to activate the production of estrogen and testosterone. Inhibition of the hypothalamo-pituitary gonadal (HPG)-axis leads to hypoestrogenia and, when combined with testosterone treatment, can mimic current clinical treatment for gender-affirming therapy in adolescents. The majority of GAHT research has revolved around cardiovascular, endocrine, and metabolic health (12). Although limited, the majority of studies investigating musculoskeletal health with GAHT treatment have focused on changes in bone health (12–14). However, connective tissues like tendon are critical for skeletal growth and mobility. Therefore, a need exists to understand the potential effects of pubertal suppression on functional properties of connective tissues, like tendon, to provide guidance on therapeutic intervention, as well as training and injury recovery, for adolescent and young adult transgender and gender diverse patients.

Previously, Dela Cruz et al. showed that GnRHa-implanted animals had reduced levels of luteinizing hormone (LH) and estradiol and decreased uterine and ovarian weights. T only and GnRHa + T treated animals also had sustained elevated levels of testosterone, suppressed LH levels compared to controls. Paired ovarian weights were reduced in the T only and GnRHa + T groups compared with the control and GnRHa-only groups (15, 16) thus proving the validity of this as a mouse model of transmasculine gender affirming care.

We and others have recently reported the effects of transmasculine gender affirming hormone treatment on skeletal growth in which puberty suppression with GnRHa results in longer femurs and testosterone treatment results in shorter femurs in female-born mice (13, 17). In this study, we examined how tendon function is influenced by GnRHa treatment, with and without subsequent testosterone administration during peripubertal growth in female-born mice. Specifically, we measured tendon structure and function in adult female-born mice subjected to pubertal suppression followed by testosterone treatment by assessing changes in mechanical properties, cell density and collagen structure and alignment.

## MATERIALS AND METHODS

### Experimental Design

All procedures were approved by the Institutional Animal Care and Use Committees at the University of Michigan. Mice were obtained post-euthanasia following tissue harvests for *in vitro* fertilization (IVF) (15) and femur characteristics (13). C57BL/6N female-born mice (n=26) and age-matched male-born mice (n= 6) were used in this study. Female-born mice were assigned to one of four experimental groups to mimic gender-affirming hormone therapy (GAHT) in human adolescents: Sham only (sham surgery with empty silastic tubing implant throughout the duration of the experiment; n=7), GnRHa only (for pubertal suppression only; n=6), Testosterone (T) only treatment (for masculinizing transition only; n=5), and GnRHa + T treatment (combined pubertal suppression and masculinizing transition; n=5). Treatment was initiated at P26, which is prior to when female-born mice enter sexual maturity. For GnRHa treated groups, mice were implanted with GnRHa subcutaneously (Goserelin acetate implant, 3.6mg, ZoladexRV, Astra Zeneca, UK). Control and T-only groups underwent a sham surgery with no implantation. After 3 weeks post-implantation, mice were subcutaneously implanted with either testosterone (T, 10mg dissolved in ethanol) in silastic tubing or with empty silastic tubing. Six weeks after T implantation (P89), mice were either euthanized or a subgroup was established to study the effects of T cessation (Washout subgroups). For these subgroup, the T-treated silastic tubing was removed and mice were euthanized two weeks later followed by ovary isolation and IVF experiments (extended sub-group, published in Dela Cruz et al. (15)). Immediately after euthanasia for all groups, mice were weighed, and hindlimbs were dissected for either histology or biomechanical testing (biomechanics, n=1 Sham + T group, n =2 GnRHa + T group; represented on all graphs as open circles; histology n=3/group) (Fig. 1). An additional group of naïve C57BL6N male and female mice were euthanized at P26 and the terminal experimental time point (P89), and Achilles tendons were dissected for RNA isolation and quantitative polymerase chain reaction (qPCR) to compare tendon-associated gene expression between male and female mice.

**Figure 1.**
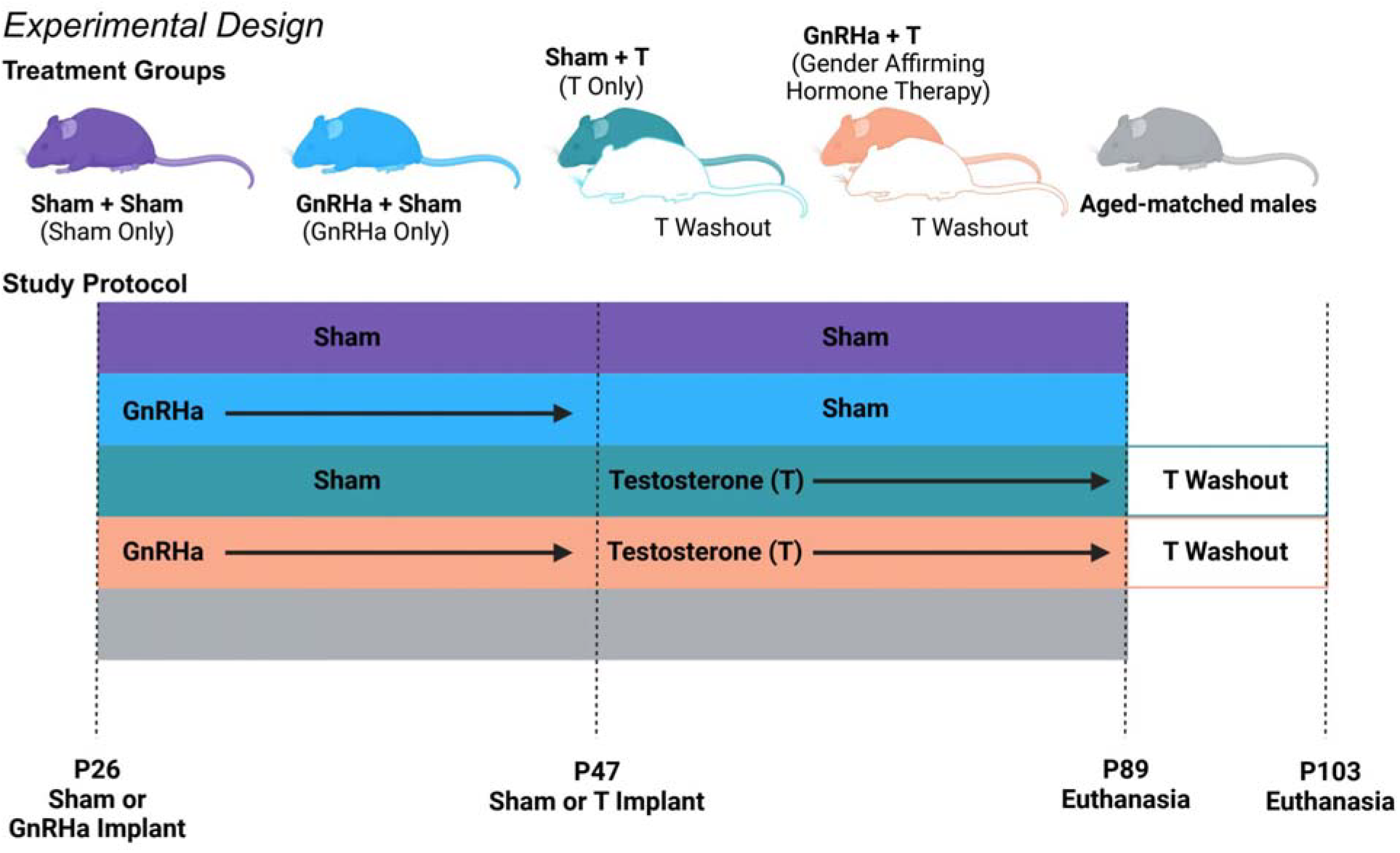
Summary of experimental design.

For the GnRHa + Sham (n=6) and GnRHa + T (n=5) groups, mice were subcutaneously implanted with GnRHa (3.6mg) at P26. Control (n=7) and T-only (n=5) groups underwent a sham surgery with no implantation. At 3 weeks post-implantation or sham surgery, mice were subcutaneously implanted with silastic tubing containing testosterone (10 mg) or empty silastic tubing (control and GnRHa + Sham group). Six weeks after T-implantation, mice were either euthanized or had a removal of their T implant. After a two-week washout period all remaining mice were euthanized.

### Biomechanical testing of Achilles tendon

Prior to biomechanical testing, carcasses were thawed at 4°C (<24 hours before testing) and prepared for biomechanical testing and photogrammetry. Briefly, the plantaris tendon and muscle were carefully removed and the Achilles tendons and calcaneus were dissected intact. Cross-sectional area (CSA) measurements were measured using photogrammetry with a custom pinch clamp holder attached to a motor controller (Arduino, Ivrea, Italy). Fifty consecutive images of the tendon-bone complex were acquired at 9° rotation steps using a macro lens (Basler Fujinon Lens, Ahrensburg, Germany). Images were converted from sparse to dense point clouds, and an STL surface mesh of the tendon was generated using Metashape software (Agisoft, St. Petersburg, Russia). CSA was measured from a slice analysis tool in Dragonfly (Comet Technologies Canada Inc., Montreal, Canada) using the smallest area within the Achilles tendon.

A custom 3D printed fixture was used to secure the calcaneus (FormLabs 3B, Somerville, MA, USA), and the proximal Achilles tendon was clamped in a thin film grip (Imada, Northbrook, IL, USA). The assembled tendon-grip unit was placed in a custom phosphate buffered solution (PBS) bath maintained at 37°C via a temperature controller (MA160, Biomomentum Inc., Laval, Quebec, Canada) and secured with a pin to a tensile testing frame with a multi-axis load cell (±70 N; Mach-1 VS500CST, Biomomentum, Laval, Quebec, Canada). Samples were preloaded to 0.1 N, and gauge length (*L*_0_) was measured (4.01 ± 0.17 mm) as the distance between the calcaneal fixture and the thin film grip.

Tendons were preconditioned for 10 cycles (±0.05 N at 0.03 Hz) followed by load to failure at 0.1 mm/min. Load and torque data (six degrees of freedom) were collected to assess off-axis loading for the duration of the experiment. Using Mach-1 Analysis and custom R script (v4.2.2 or later, The R Project for Statistical Computing, Vienna, Austria), the mechanical properties of the Achilles were calculated from force-displacement data. Engineering stress was calculated as the instantaneous force divided by the original CSA (calculated from photogrammetry). Instantaneous grip-to-grip strain was calculated as the displacement divided by the original gauge length, *L*_0_. Maximum load, maximum stress, and maximum strain were calculated using R. Stiffness and tangent modulus were determined from load-displacement and stress-strain data respectively, using the piecewise linear segmentation by dynamic reprogramming recursion package (dpseg) in R. Data from female-born mice were compared using a one-way ANOVAs and corrected for multiple comparisons using Tukey’s multiple comparisons tests (Prism v9.0+; Graphpad, LaJolla, CA, USA).

### Histology Preparation and Imaging

Distal hindlimbs from GAHT treatment and control mice (n = 3 per group) were decalcified in 14% ethylenediaminetetraacetic acid (EDTA) and processed for paraffin embedding. Tissues were sectioned at 5 μm in the sagittal plane and stained with Hematoxylin and Eosin (H&E) or Picrosirius Red (PSR) (Stat Labs, McKinney, TX, USA), and cover-slipped with an acrylic mounting media (Shandon Mount, ThermoFisher, Waltham, MA, USA). H&E slides were imaged on a bright-field microscope (ECLIPSE Ni-U, Nikon) at 10x and analyzed using QuPath (https://qupath.github.io). PSR slides were imaged with a 10x objective on an epiflourescent microscope (dmi600b, Leica) and analyzed using quantitative polarized light imaging (qPLI) analysis (ThorLabs camera)(18).

### QuPath Analysis

H&E slides were imaged and collected for cell density analysis by QuPath, an open-source software for digital pathology. First, enthesis and insertional tendon areas were selected using morphological features. Next, color deconvolution was performed by QuPath to digitally separate stains within each tissue sample. Positive cell detection was then performed to differentiate cells. For H&E staining, hematoxylin positive nuclei were identified, and cell number and cell density within each tissue sample were collected as a result. QuPath cell density analysis was performed by two reviewers and all cell and area counts were averaged. Enthesis and insertional area and cell density from female-born mice were compared using a one-way ANOVAs and corrected for multiple comparisons using Tukey’s multiple comparisons tests (Prism v9.0+; Graphpad, LaJolla, CA, USA). Cell number and area correlation plots were analyzed using simple linear regression and Pearson’s correlation tests (Prism v9.0+; Graphpad, LaJolla, CA, USA).

### qPLI Analysis

The degree of linear polarization (DoLP) and angle of linear polarization (AoLP) images were acquired using a polarization camera (Thorlabs, Newton, New Jersey, USA) and a circular polarizing lens (Edmund Optics, Barrington, New Jersey, USA). The mean DoLP and standard deviation of the AoLP were analyzed using Math and SciPy Stats libraries in Python (v3.12.1, Python, Wilmington, Delaware, USA). Data from female-born mice were compared using a one-way ANOVAs and corrected for multiple comparisons using Tukey’s multiple comparisons tests (Prism v9.0+; Graphpad, LaJolla, CA, USA).

### RNA isolation and qRT-PCR

We verified gene expression of enthesis and tendon reference genes: *Tnc, Tnmd, Scx*, and *Sox9* (Integrated DNA Technologies) with qRT-PCR. Achilles tendons from aged-matched female-born and male-born mice at P26 and P89 (*n* = 4/group) were immediately dissected under ribonuclease (RNase)–free conditions and snap-frozen. Tissues were mechanically pulverized in TRIzol, and total RNA was isolated using spin-columns (PureLink RNA mini kit, Thermo Fisher Scientific) with on column genomic DNA digestion (RNase-free DNase, QIAGEN). The quality and quantity of RNA were checked with a NanoDrop spectrophotometer (Thermo Fisher Scientific). RNA with a 260/280 ratio graeter than 2.0 was reverse-transcribed to cDNA (SuperScript IV VILO Master Mix, Thermo Fisher Scientific). After conversion, 10 ng/uL of cDNA was used per reaction. Quantitative real-time reverse transcription polymerase chain reaction (qRT-PCR) was performed using a CFX96 Real-Time PCR Detection System (Bio-Rad) with Power SYBR Green PCR Master Mix (Thermo Fisher Scientific). *Rplpo* was used as the reference gene. Primer information is provided in Table 1. Data were compared using two-way ANOVAs and corrected for multiple comparisons with Tukey’s multiple comparisons tests (Prism v9.0+; Graphpad, LaJolla, CA, USA).

**Table 1.** qPCR primer information. IDT Predesigned qPCR Assays: intercalating dyes, primers only.

| Assay ID | Gene Query | NCBI Gene Symbol | Ref Seq # | Detects All Variants | Exon Locations |
| --- | --- | --- | --- | --- | --- |
| Mm.PT.58.43894205 | Rplp0 | Rplp0 | NM_007475(1) | Yes | 5-6 |
| Mm.PT.58.8090418 | Tnc | Tnc | NM_011607(1) | Yes | 18-19 |
| Mm.PT.58.13530921 | Tnmd | Tnmd | NM_022322(1) | Yes | 5-6 |
| Mm.PT.58.31750069 | Scx | Scx | NM_198885(1) | Yes | 1-2 |
| Mm.PT.58.42739087 | Sox9 | Sox9 | NM_011448(1) | Yes | 1-2 |

## RESULTS

### Achilles tendons in female-born mice treated with gender-affirming hormone therapy were stronger

Treatment with T only led to significantly increased body weight of female-born mice (Fig. 2A). We did not find any significant body weight differences in groups treated with GnRHa with or without T (Fig. 2A). To investigate the effect of GAHT on tendon function, Achilles tendons were uniaxially tested after euthanasia at either P89 or P103. We found that all GAHT treatment strategies resulted in an increase in load at failure (max load) compared to sham-treated controls (Fig. 2B). Additionally, treatment with T increased variation in stiffness (Fig 2C). No significant differences in CSA were found between any groups (Fig. 2C). However, we found that only treatments with GnRHa, with and without T, significantly increased the stress at failure (max stress, Fig. 2D&E). Finally, we did not identify any significant differences between groups related to tangent modulus (Fig. 2F).

**Figure 2.**
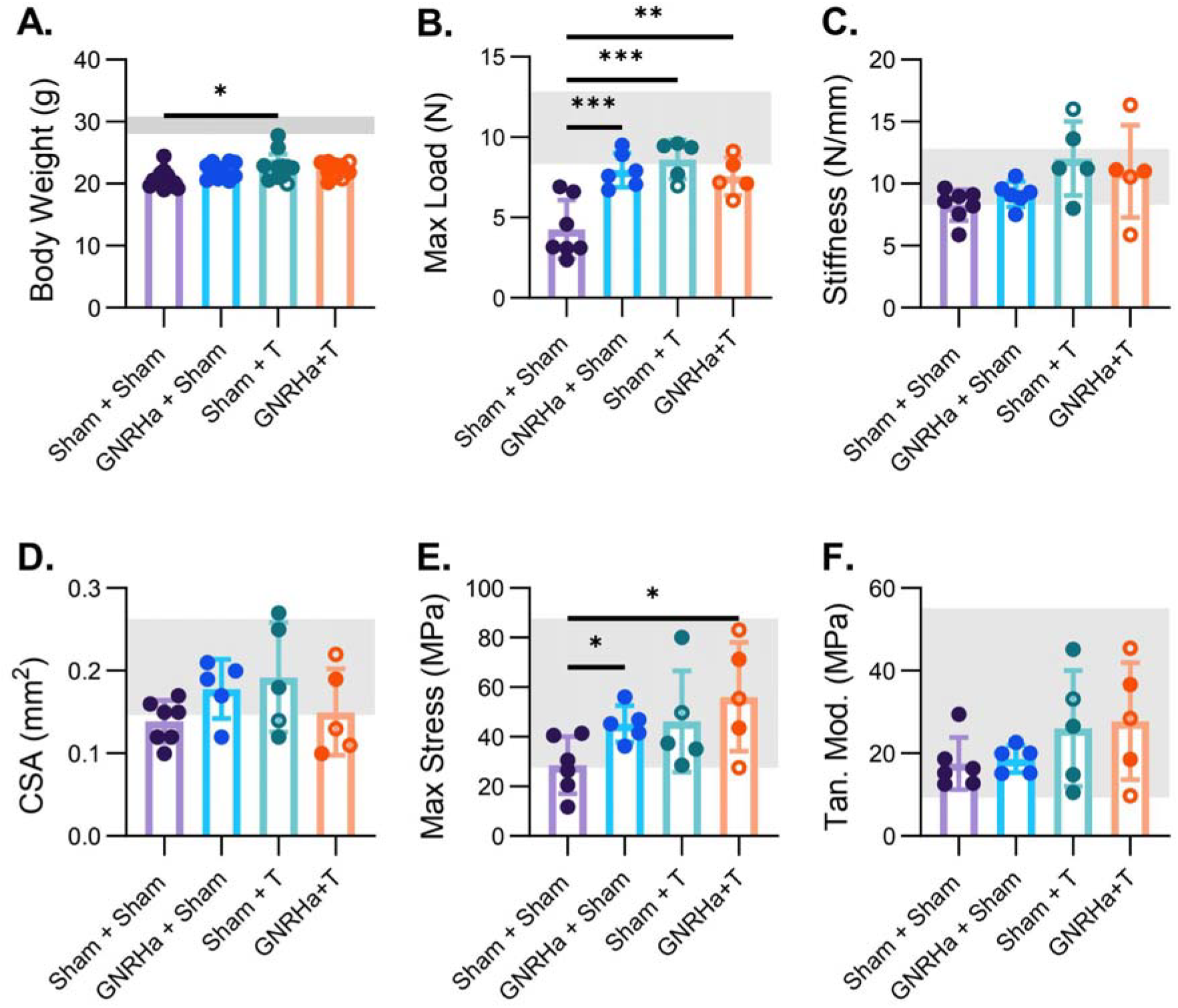
Mechanical testing of Achilles tendons showed maximum load of tendons was increased in all treatment groups compared to controls while GnRHa treatment with and without testosterone increased maximum stress in tendon. Treatment with T significantly increased the body weight (g) of female-born mice after treatment (A). Maximum load (N) of Achilles tendons was significantly increased in all groups compared to controls (B). Achilles tendon stiffness remained unchanged between all groups (C). Cross sectional area (CSA) (mm) of Achilles tendons remained unchanged between all groups (D). Maximum stress (MPa) of Achilles tendons was significantly increased in GnRHa treated mice with or without testosterone compared to controls (E). Tangent Modulus (MPa) of Achilles tendons remained unchanged between all groups (F). Data are presented as biological replicates (individual dots) and mean +/-standard deviation. Mice who were given a two-week T washout are represented as open circles. Gray horizontal bars represent range for age-matched male control mice, n=3. * p< 0.05; ** p< 0.01; ***p<0.001.

### Female-born mice treated with GnRHa had increased enthesis cellular density and no change to collagen organization

GnRHa treatment alone led to significantly increased enthesis cell number and density compared to all other groups (Fig 2A&B). We found no significant differences between enthesis area between all groups (Fig. 2C). We also found that there was not a strong correlation between area of the enthesis and cell number when comparing all groups (r^2^=0.329; p = 0.0510 Fig. 2D). Furthermore, we also saw no change in cell number or density in any treatment group in the insertional tendon and there was not a strong correlation between area of the insertional tendon and cell number when comparing all groups (r^2^=0.1402; p = 0.2305) (Fig. 2I-L). We found that collagen organization of the enthesis and insertional tendon was also not affected by any GAHT treatment strategy (Supplemental Fig. 1).

### Tendon and enthesis markers were age dependent in both pre- and post-pubertal mice and *Tnmd* was sex dependent in pre-pubertal mice

To identify if tendon and enthesis markers differ between sexes (without GAHT treatment) at key time points used in this study, we measured relative levels of *Scx, Tnc, Tnmd*, and *Sox9* at P26 (start point of GAHT) and P89 (end point of GAHT). We found that gene expression of tendon and enthesis markers, *Scx, Tnc* and *Tnmd*, were age dependent (Fig. 4 A,C&D). Relative gene expression at P89 in the male born mice was reduced for *Tnc* and *Tnmd* compared to male mice at P26 (Fig.4C&D). Additionally, we saw no change in the relative expression of *Sox9* regardless of sex or age (Fig. 4B). Finally, *Tnmd* was the only marker we selected that was significantly different between male and female-born mice, and female-born mice had a reduced expression of *Tnmd* at P26 compared to male mice (Fig. 4D).

## DISCUSSION

The potential effects of pubertal suppression on the functional properties of musculoskeletal tissues, like tendon, is an important and understudied topic. In tendon, the effects of estrogen and T have recently been reported to have similar effects but have differing mechanistic targets during healing (19). RNA sequencing perfomed on the supraspinatus tendon 2 weeks post laceration suggested these hormones can mediate gene expression during the healing process (19). These considerations for the differences between sex hormones and the impact on tendon health are important for transgender and gender diverse youth who may use GnRHa to delay or inhibit secondary sex characteristics.

The potential effects of the role of sex hormones on tendon functionality has not been explored extensively in pre-pubescent populations, and the data within the adult population is limited in scope and yields conflicting results. Previous studies have shown treatment with hormones following disease or injury has a positive effect on tendon functionality (20–22). For example, treatment with estradiol increased tensile strength, stiffness and maximum load in Achilles tendinitis in a rat model (20). However, in non-diseased tendons estrogen has been shown to influence tendon morphology and limit tendon functionality. Elevation of estradiol levels through oral estradiol replacement therapy (ERT) resulted in increased collagen synthesis and reduced fibril size in postmenopausal female-born patients compared with female-born patients exposed to low-estradiol concentration. However, ERT therapy reduced stiffness in postmenopausal female-born patient possibly due to immature crosslinking from increased synthesis (21). Treatment with anabolic androgenic steroids treatment strongly decreased matrix metalloproteinase-2, a known regulator of collagen remodeling, concentration in the three regions of the superficial flexor tendons (22). Interestingly, we found that hormone suppression with GnRHa improved tendon functionality with or without testosterone in female-born mice. While treatment with testosterone alone increased the maximum load of the tendon, maximum stress remained the same as female-born controls when accounting for changes to CSA. Additionally, we saw no differences in overall alignment of the tendon regardless of treatment (Fig. 2). These data taken together brings to question the role of estrogen and testosterone on factors that influence overall collagen deposition and remodeling that could lead to improved tendon functionality.

Although the effects of the administration of sex hormones as a regulatory factor in the metabolism of cartilage, bone and muscle have been studied in depth, little is known about the role of sex hormones in tendon and enthesis (23). Ovariectomized mice exhibit increased chondrocyte proliferation compared with sham controls, and these results were negated in estrogen receptor beta knockout mice with ovariectomy (24). These findings suggest estrogen is a negative regulator for chondrocyte proliferation. These data support our findings that hormone suppression with GnRHa significantly increased cell density in the female-born enthesis, a fibrocartilage tissue. However, we also found that puberty suppression with GnRHa, when supplemented with testosterone, did not affect cell density compared to controls. In tendon derived cells from male-born rats, treatment with estrogen and estrogen-like analogues significantly increased cell proliferation *in vitro* while estrogen agonists only decreased proliferation when given in the presence of estrogen (25). We found that hormone suppression with GnRHa with and without testosterone as well as testosterone alone had no effect on the cellular density of the insertional tendon (Fig. 3). The differences in findings between ours and previous studies could be explained by lower levels of circulating estrogen in the Achilles tendon than the doses given *in vitro*.

**Figure 3.**
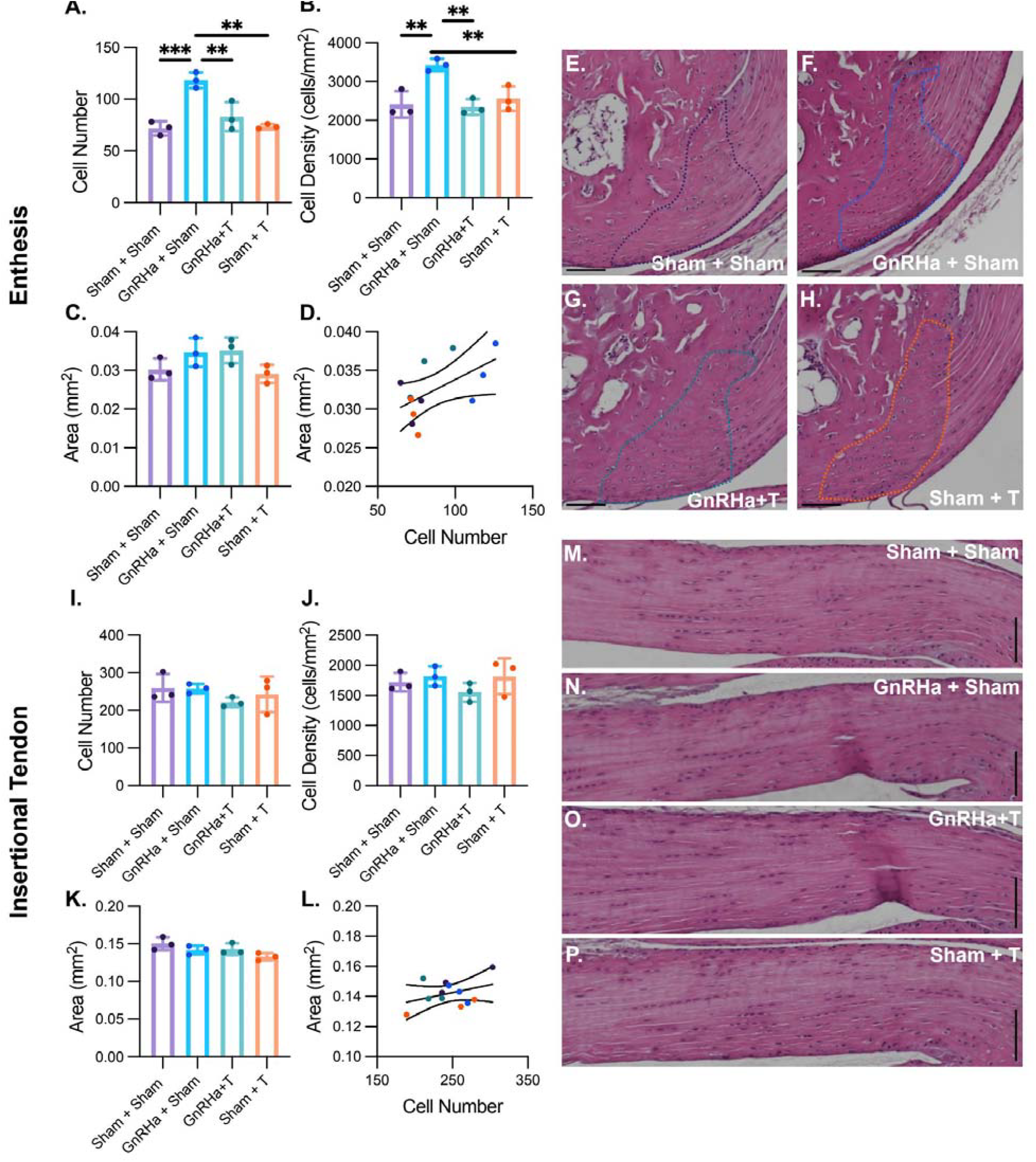
GnRHa significantly increased enthesis cellular number and density. Treatment with GnRHa significantly increased Achilles enthesis cell number compared to all groups (A). Treatment with GnRHa significantly increased Achilles enthesis cell density (cells/mm^2^) compared to controls and GnRHa + T treated female-born mice (B). Achilles enthesis area (mm^2^) remained unchanged between all groups. (C) Linear regression of Achilles enthesis area and cell number between all groups, r^2^=0.329 (D). Representative images of Achilles enthesis areas used for cell number and density calculations (E-H). Achilles insertional tendon cell number remained unchanged between all groups. (I) Achilles insertional tendon cell density (cells/mm^2^) remained unchanged between all groups. (J). Achilles insertional tendon area (mm^2^) remained unchanged between all groups. (K) Linear regression of Achilles insertional tendon and cell number and area between all groups, r^2^=0.1402 (L). Representative images of Achilles insertional tendon used for cell number and density calculations (M-P). Data are presented as biological replicates (individual dots) and mean +/-standard deviation. n=3. ** p< 0.01; ***p<0.001.

We observed in male and female-born pre-pubertal mice only the tendon maker *Tnmd* was sex dependent (Fig.4). It has been shown that loss of *Tnmd* in mice does not produce severe developmental phenotypes. However, *Tnmd* deficient mice do exhibit reduced tendon cell density, reduced tenocyte proliferation, and elevated collagen fibril size (26). Additionally, *Tnmd* may also play a similar role in the enthesis. In mouse enthesis mesenchymal cells, single cell RNA sequencing defined a subpopulation termed as “enthesoblasts,” which had the most enriched profiles of matrix deposition and tissue development. These cells had high expression of tendon and chondrogenic markers which included *Tnmd* (27). has been shown that *Tnmd* may be negatively regulated by estrogen like compounds, *Tnmd* expression was increased in tendons in female-born rats who underwent an ovariectomy compared to intact tendons. While treatment with genistein, a plant-derived, naturally occurring isoflavone phytoestrogen, returned values comparable to controls (28). Taken together, we believe an increase in *Tnmd* could be a possible mechanistic target responsible for the increase in tendon functionality in response to hormone suppression with GnRHa with and without testosterone. Future studies should explore how changes in *Tnmd* are associated with changes to testosterone and pubertal suppression in the context of tendon health.

**Figure 4.**
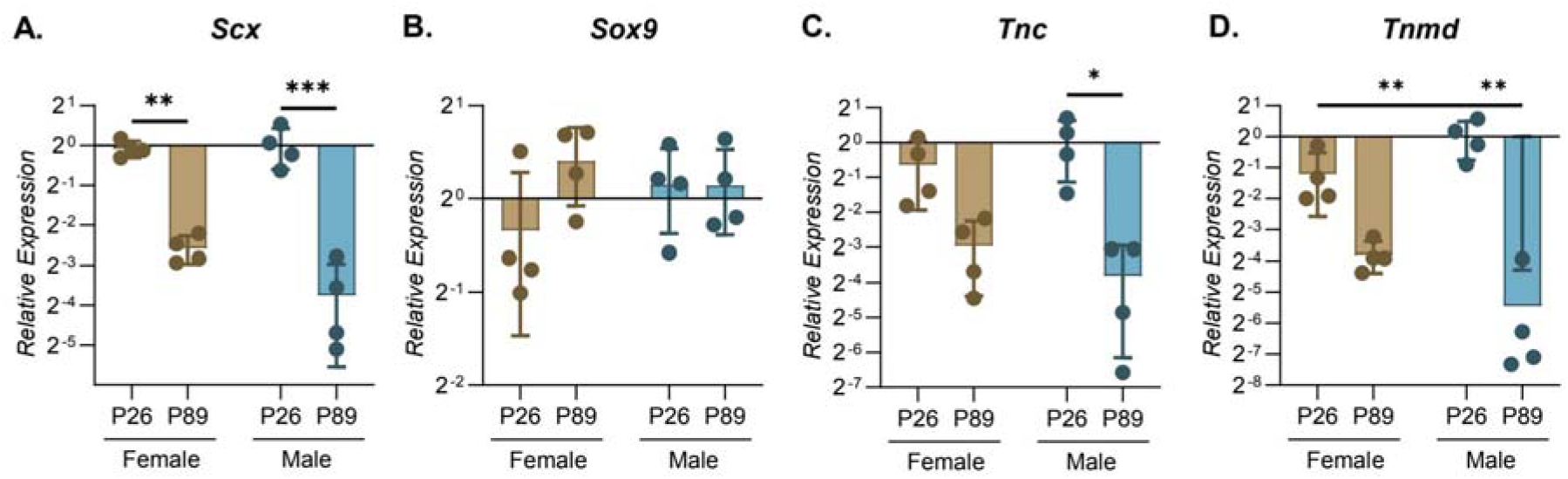
Tendon and enthesis markers *Scx* and *Tnc* are age dependent while *Tnmd* is age- and sex-dependent in prepubertal mice. Achilles tendon *Scx* relative gene expression was significantly reduced in female and male born mice at P89. (A) Achilles tendon *Sox9* relative gene expression had no change in female and male born mice at P26 and P89. (B) Achilles tendon *Tnc* relative gene expression trended down in female-born and was significantly reduced in male born mice at P89 compared to P26. (C) Achilles tendon *Tnmd* relative gene expression was significantly reduced in female-born mice compared to male mice at P26 and was significantly reduced at P89. (D) Data are presented as biological replicates (individual dots) and mean +/-standard deviation. n=3. * p<0.05, ** p< 0.01; ***p<0.001.

In summary, this work demonstrates that gender affirming hormonal therapy can influence enthesis and tendon functionality in mice. While puberty suppression through GnRHa does change enthesis cellular behavior, these gender affirming therapies (e.g GnRHa + T) do not have any significant effect on the histological properties of the enthesis or tendon. Additionally, these gender affirming therapies significantly improved the overall tendon functionality of female-born mice. Taken together, these findings begin to answer important questions of the potential effects of pubertal suppression on the functional properties of the tendon and enthesis and may help to provide clinical guidance and training perspectives for transgender and gender diverse patients and athletes.

## ACKNOWLEDGMENTS

Schematics designed using Biorender. We are grateful for histological support from Emma Snyder-White and Carol Whitinger (through the Michigan Integrative Musculoskeletal Health Center Core).

## DISCLAIMERS

The authors have no disclaimers.

## AUTHOR CONTRIBUTIONS

MLK, LH, and AS conceived and designed research; LH, TP, PC, SS, SB, NM, and CDC performed experiments; LH, SB, and SG analyzed data; LH, BH, and MLK interpreted results of experiments; LH prepared figures; LH and SG drafted the manuscript; LH, BH, SS, NM, and MLK edited the manuscript; ACA, MLK, MM, VP, and AS approved final version of manuscript.

