## Supplemental for "Functional Changes to Achilles Tendon and Enthesis in a Mouse Model of an Adolescent Masculine Gender-Affirming Hormone Treatment"

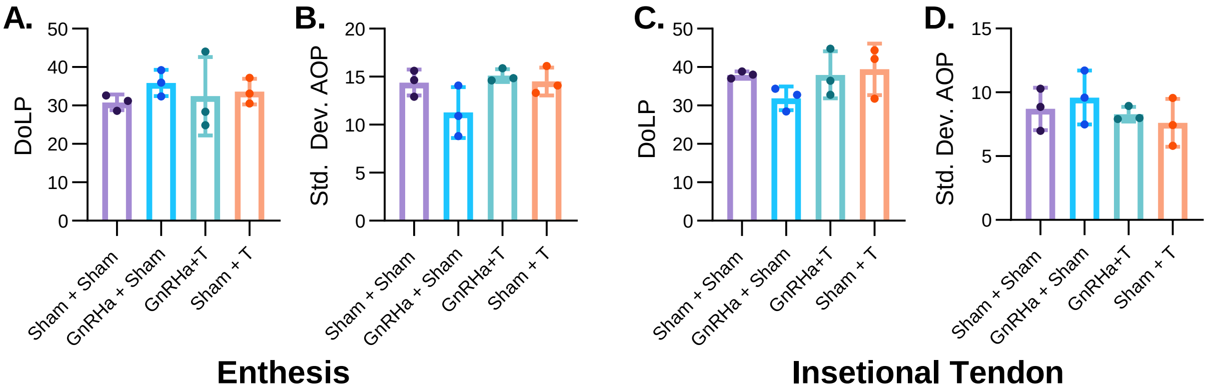


**Supplemental Figure 1:** Treatment with gender affirming hormone therapy had no efffect on tendon or enthesis organization. There were no change to the degree of linear polarization (DoLP) (A) or  the standard deviation of the angle of polarization (std. dev. AOP) (B) of the defined enthesis area between groups.There were no change to the (DoLP) (C) or the std. dev. AOP (D) of the defined insertional tendon area between groups. n=3.
